# Specification of embryonic shoot stem cells via a small RNA-driven morphogenic circuit

**DOI:** 10.64898/2026.08.21.746189

**Authors:** Qi Li, Cecilia Lara-Mondragón, Antje Feller, Arvid Herrmann, Steffen Knauer, Mary Galli, Xixi Zheng, Caroline Marcon, Frank Hochholdinger, Andrea Gallavotti, Marie Javelle, Marja C.P. Timmermans

## Abstract

The specification of embryonic stem cells capable of self-renewing and differentiation into virtually any cell type, is one of the most consequential events in the development of a multicellular organism. Yet the mechanisms establishing embryonic stem cell fate remain poorly understood, particularly in monocotyledonous cereals. Using the classic mutant *leafbladeless1-raggedseedling1*, we show that the small RNA tasiARF acts as a primary epidermis-derived morphogenic signal that organizes shoot stem cell specification in the maize embryo. tasiARF restricts expression of the AUXIN RESPONSE FACTOR 3 (ARF3) transcription factor, which modulates cell wall mechanics and guides the differential localization of PIN-FORMED (PIN) auxin efflux carriers, establishing a localized auxin minimum permissive for stem cell fate. Interestingly, loss of this auxin minimum and the associated shoot stem cell defects in tasiARF-deficient embryos are buffered by natural variation at a quantitative trait locus (QTL) controlling expression of MICROTUBULE-ASSOCIATED PROTEIN 65-3 (MAP65-3), a critical factor determining cell division orientation, which reshapes auxin dynamics and restores stem cell specification, and is itself under tasiARF-ARF3 control. Thus, embryonic shoot stem cell specification in maize is governed by an intricate morphogenic circuit that couples small RNA-mediated positional information to a self-stabilizing network interdependently linking cell wall mechanics, auxin signaling, and cell division patterning. This mechanistic framework reveals the redeployment of an ancient small RNA pathway as a lineage-specific innovation to establish a conserved, stem cell-permissive low auxin environment within the divergent embryonic architecture of monocotyledonous cereals. More broadly, it identifies molecular entry points for the engineering of embryogenic competence and regeneration capacity for the improvement of cereal crops.

**In brief:** The small RNA tasiARF initiates a self-reinforcing morphogenic circuit connecting cell wall mechanics, auxin signaling, and regulated cell division to specify embryonic shoot stem cell fate, revealing how lineage-specific regulatory innovation can preserve a conserved developmental output across divergent embryonic architectures.

**Highlights:**

- The small RNA tasiARF is an epidermal morphogenic signal specifying embryonic stem cell fate in maize
- tasiARF signaling creates a local stem cell-permissive mechanical and auxin environment
- Natural variation influencing cell division patterning buffers embryonic shoot stem cell specification
- Embryonic stem cells arise through lineage-specific regulatory innovation adapted to embryonic architecture

## Introduction

Embryonic stem cells are a population of pluripotent cells capable of self-renewal and differentiation into virtually any cell type of a multicellular organism^1–3^. Their specification is one of the most consequential events in development. In both animals and plants, a stem cell population must be singled out from an initially equivalent group of cells, organized into a functional niche, and shielded from the differentiation pressures acting on surrounding tissues, all while the embryo undergoes rapid morphogenesis. How a small subset of embryonic cells acquires stem cell identity remains a central question in developmental biology, with far-reaching implications for both medicine and agriculture^4–7^.

In plants, embryonic stem cells are established within the shoot and root apical meristems, specialized niches that maintain stem cell identity and coordinate the formation of new organs throughout the plant’s lifetime^1,2^. In the model eudicot *Arabidopsis thaliana*, early embryogenesis proceeds through a stereotyped sequence of cell divisions coupled with progressive fate transitions that culminates in the establishment of the shoot stem cell niche at the embryonic apex^8,9^. In this process, the patterned distribution of the phytohormone auxin plays a central role. Polar auxin transport concentrates auxin into two maxima at which the cotyledons emerge, thereby depleting auxin from the boundary region between them. This local auxin minimum provides a permissive environment for shoot stem cell specification and defines the position where the shoot meristem will form. Embryonic shoot stem cell fate is therefore coupled to the establishment of bilateral symmetry, linking cotyledon patterning directly to stem cell niche formation.

In monocotyledonous cereals such as maize (*Zea mays*), early embryogenesis follows a markedly different developmental strategy, fundamentally changing this geometry^9–11^. The embryo undergoes relatively stochastic divisions to form a club-shaped proembryo that abruptly breaks radial symmetry to establish dorsoventral or adaxial-abaxial polarity. At this stage, shoot stem cell precursors, distinguished by increased proliferation and elevated cytoplasmic density, are specified on the dorsal/adaxial side opposite the single cotyledon (scutellum)^9–11^. The transition to a monocotyledonous architecture therefore fundamentally reshaped the spatial context within which shoot stem cells arise, raising the question whether stem cell fate in cereals likewise requires a low auxin environment, how such a domain would be established, and to what extent the underlying principles are conserved. Addressing these questions is essential for understanding how embryonic stem cell specification is achieved and has diversified during plant evolution, while also opening new avenues for crop improvement through engineering embryogenic competence and regeneration potential^5–7^.

Here, we show that the conserved 21-nt small regulatory RNA tasiARF acts as an epidermis-derived positional signal that coordinates local patterning to specify the embryonic shoot stem cell niche in maize. tasiARF spatially restricts expression of the AUXIN RESPONSE FACTOR3 (ARF3) transcription factor, thereby modulating local cell wall and mechanical properties and establishing a low auxin environment permissive to stem cell specification. In tasiARF-deficient embryos, a local auxin minimum fails to form, causing a failure of niche formation; a defect rescued by natural variation at a QTL regulating expression of MAP65-3, a critical factor determining cell division orientation and cell plate formation. This QTL reshapes auxin dynamics and restores stem cell specification, and is itself controlled by the tasiARF-ARF3 regulatory module. These findings show that embryonic shoot stem cell specification in maize is driven by a self-stabilizing morphogenic circuit initiated by a conserved small RNA signal, that connects cell wall mechanics and auxin signaling with cell division patterning to confer robustness to the developmental program. They reveal how lineage-specific regulatory innovation can preserve a conserved developmental output across divergent embryonic architectures.

## Results

### tasiARF as an adaxial protodermal signal specifies the shoot stem cell niche

Trans-acting small interfering RNAs (tasiRNAs) are a class of plant-specific small RNAs that function as key morphogenic signals guiding plant development^12,13^. Like microRNAs, tasiRNAs act non-cell-autonomously providing positional information that patterns gene expression at the post-transcriptional level^14–16^. Mutants defective in tasiRNA biogenesis in monocots frequently show defects in stem cell homeostasis, and in the most extreme case fail to specify a functional shoot stem cell niche during embryogenesis (**Supplementary Fig. 1**)^17–20^. These mutants therefore provide a unique opportunity to identify conserved principles and lineage-specific innovations in embryonic stem cell specification in monocot cereals.

In maize, the fertilized zygote divides asymmetrically to distinguish a small apical cell that undergoes a series of seemingly random divisions generating a club-shaped proembryo^9–11^. At the early transition stage (6-7 days after pollination), changes in the pattern of cell division identify a shift from radial symmetry to adaxial-abaxial polarity. At this time, the single cotyledon or scutellum begins to form, and a small cluster of cells emerges from the adaxial surface of the embryo, marking the presumptive shoot stem cell niche (**Fig. 1A-1B**). Shortly thereafter, the ring-shaped primordium of the coleoptile becomes evident and gradually enlarges to envelop the shoot stem cell domain^10–11^. At the proembryo stage, *leafbladeless1-ragged seedling1* (*lbl1-rgd1*) mutant embryos, which carry a mutation in the maize homolog of *Arabidopsis SUPPRESSOR OF GENE SILENCING 3* (*SGS3*) required for tasiRNA biogenesis^18^, are indistinguishable from wild-type. During the transition stage, however, *lbl1-rgd1* embryos develop a rounded scutellum and fail to establish both the shoot stem cell niche and the surrounding coleoptile on the adaxial side of the embryo (**Fig. 1A-1B**). As a result, *lbl1-rgd1* mutants develop a root system but lack a shoot (**Supplementary Fig. 1A**). Consistent with the perturbed specification of shoot stem cells, expression of the homeobox gene *Knotted1* (*Kn1*), a key regulator of shoot stem cell identity^21^, is markedly changed in *lbl1-rgd1* (**Fig. 1B**). In wild-type embryos, *Kn1* marks the stem cell precursors at the early transition stage, albeit not in the protoderm, and expression subsequently expands along the developing shoot-root embryonic axis^21^. In *lbl1-rgd1* mutants, *Kn1* expression remains detectable at the transition and coleoptile stages, but the domain of expression is notably reduced, with *Kn1* transcripts undetected in the adaxial protoderm as well as the immediate underlying three to four cell layers. This change in expression highlights a critical role for the tasiRNA pathway not in activating *Kn1* expression per se, but in establishing its correct spatial pattern.

**Figure 1.**
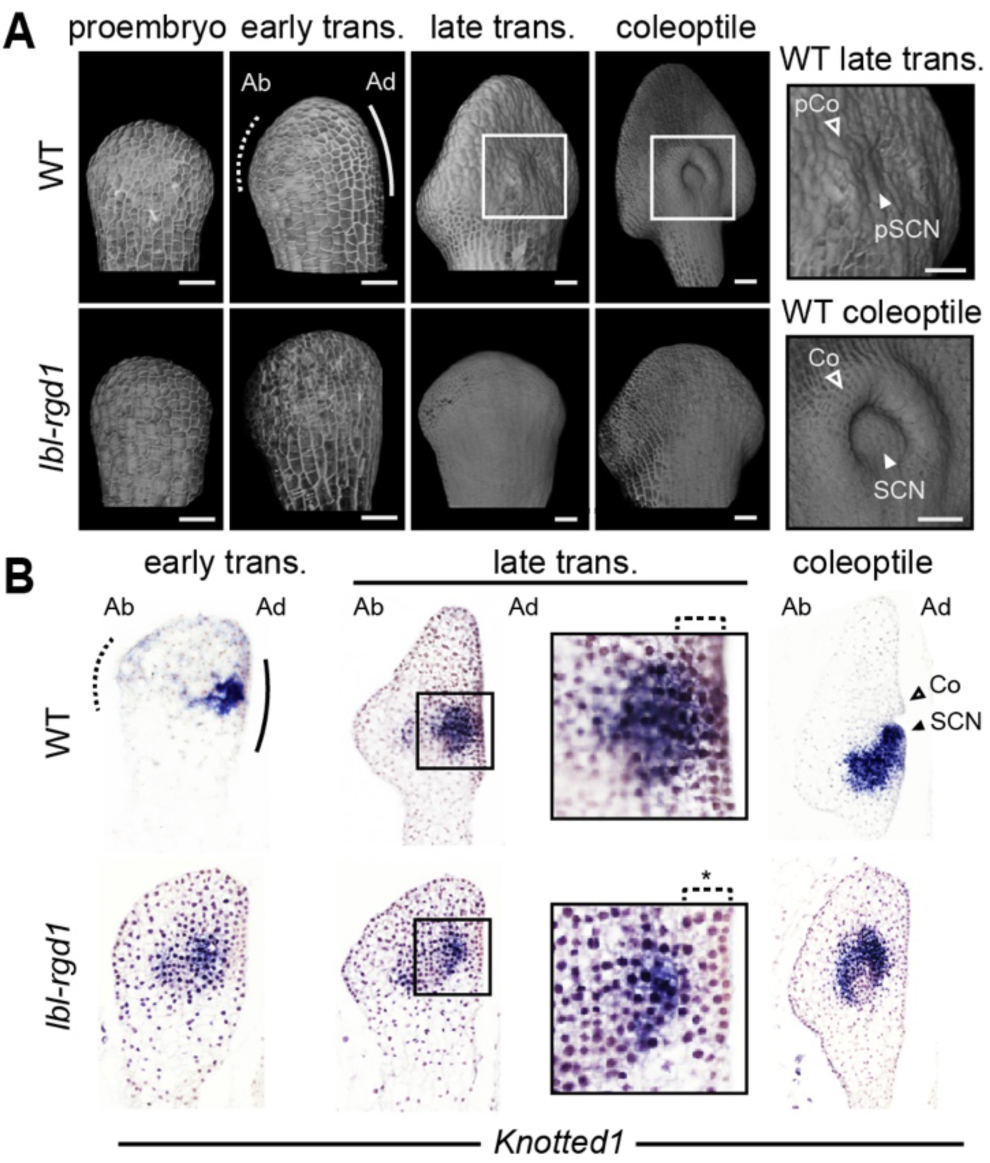
The small RNA tasiARF is required for embryonic shoot stem cell specification. (**A**) Three-dimensional reconstructions of wild-type (WT) and *lbl1-rgd1* embryos at successive developmental stages shows that adaxial-abaxial polarity becomes apparent during progression from the proembryo to early transition stage in both genotypes. In wild-type embryos, the shoot stem cell niche and surrounding coleoptile are subsequently specified on the adaxial side, whereas this patterning is perturbed in *lbl1-rgd1*. Enlarged views of the developing shoot stem cell region are shown on the right. (**B**) RNA in situ hybridization showing *Knotted1* (*Kn1*) expression along the embryonic shoot-root axis during wild-type embryogenesis. In *lbl1-rgd1, Kn1* expression is depleted from the adaxial-most cell layers. Enlarged views of the stem cell region in late transition-stage embryos highlight the spatial relationship between *Kn1* expression and the adaxial protoderm. Asterisks, absence of *Kn1* expression in *lbl1-rgd1*; dashed line, abaxial (ab); solid line, adaxial (ad); trans., transition stage; pSCN, presumptive shoot stem cell niche; SCN, shoot stem cell niche; pCo, presumptive coleoptile; Co, coleoptile. Scale bars, 50 μm. Genetic background, W22.

The localized loss of *Kn1* expression led us to hypothesize that *lbl1-rgd1* interferes with a positional signal required for shoot stem cell specification from adaxial embryonic cells. The tasiRNA, tasiARF, emerged as a strong candidate, as it functions as a morphogen-like signal that regulates AUXIN RESPONSE FACTOR 3 (ARF3) expression during adaxial-abaxial leaf patterning^13–16,18^. Indeed, tasiARF accumulates strongly in the adaxial subapical protoderm in early transition stage embryos, while its target *ARF3a* exhibits a broader expression minimum that overlaps the initiation site of *Kn1-*marked shoot stem cell precursors, reflecting the non-cell-autonomous action of tasiARF. In *lbl1-rgd1,* where tasiARF is depleted^18^, *ARF3a* accumulates ectopically in the adaxial protoderm and two to three underlying cell layers (**Fig. 2A**). These findings highlight a critical role for the tasiRNA pathway in specifying the embryonic shoot stem cell niche, with tasiARF conveying positional information to define a spatially restricted *ARF3* expression minimum on the adaxial side of the early transition stage embryo necessary for shoot stem cell identity.

**Figure 2.**
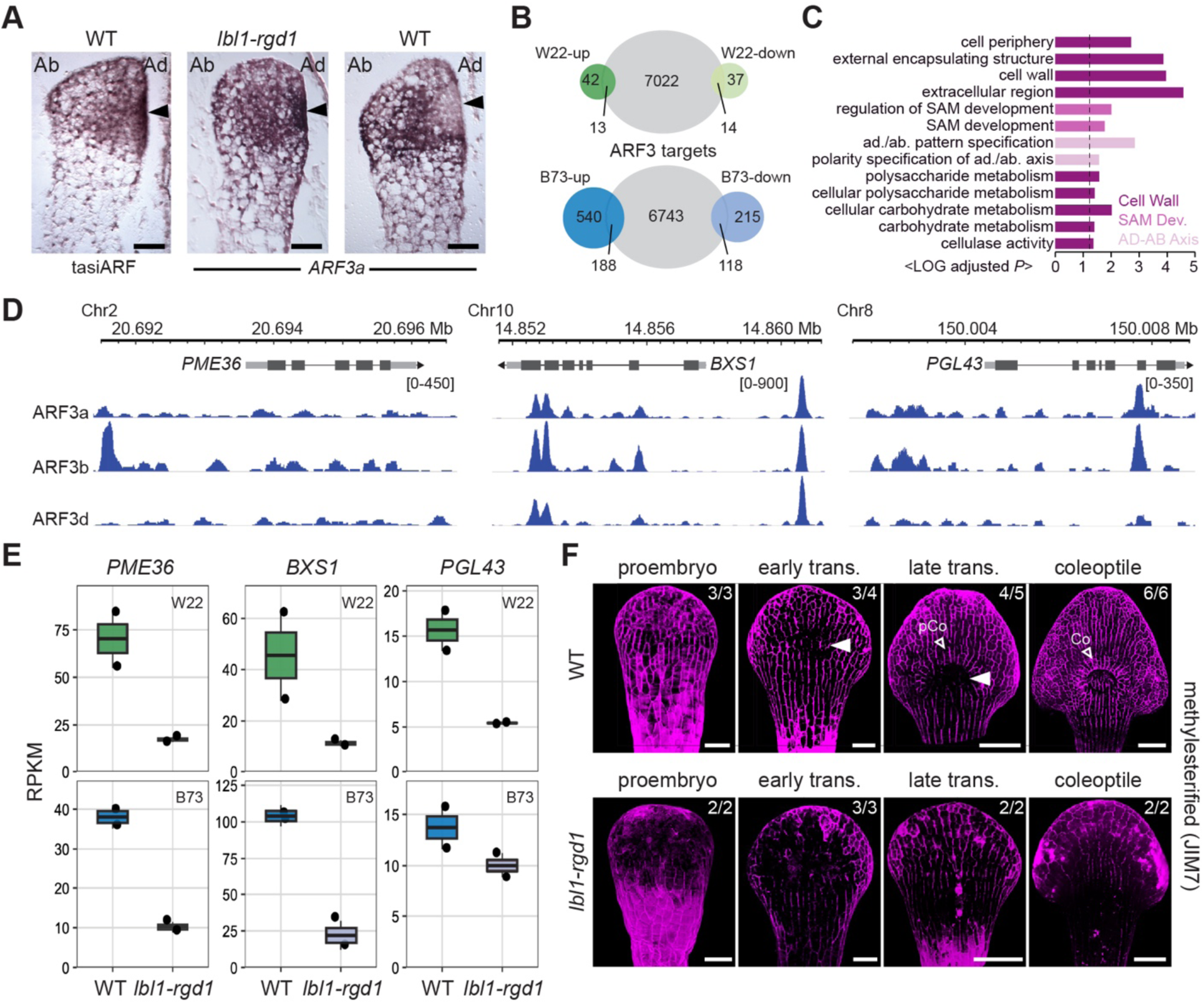
tasiARF-ARF3 signaling patterns cell wall properties during embryonic shoot stem cell specification. (**A**) RNA *in situ* hybridization of tasiARF and its target *ARF3a* in wild-type and *lbl1-rgd1* transition stage embryos shows tasiARF strongly enriched in the adaxial protoderm represses ARF3 expression in the presumptive shoot stem cell niche region. ad., adaxial; ab., abaxial. (**B**) Venn diagrams showing the overlap between ARF3 DAP-seq targets (grey; genes bound by ARF3a, ARF3b, and/or ARF3d) and genes differentially expressed in *lbl1-rgd1* early transition stage embryos in the W22 and B73 background. Up, upregulated in *lbl1-rgd1*; down, downregulated in *lbl1-rgd1*. (**C**) Gene Ontology enrichment analysis of genes in the overlaps in (B) identifies cell wall, shoot apical meristem development (SAM Dev.), and adaxial–abaxial patterning (AB–AD Axis) as enriched functional categories. (**D**) Representative DAP-seq binding profiles for ARF3a, ARF3b, and ARF3d showing occupancy in promoter and/or genic regions of the cell wall-associated genes *PME36*, *BXS1*, and *PGL43*. (**E**) RNA-seq analysis shows *PME36*, *BXS1*, and *PGL43* expression is reduced in *lbl1-rgd1* early transition stage embryos. RPKM, reads per kilobase of transcript per million reads mapped. (**F**) Adaxial views of JIM7 whole-mount immunofluorescence-stained embryos show a localized depletion of highly methyl-esterified pectin at the initiating shoot stem cell niche in wild-type transition stage embryos (arrowheads) that contrasts to the dispersed depletion of JIM7 signal in *lbl1-rgd1* sibling embryos. pCo, presumptive coleoptile; Co, coleoptile; trans., transition stage. Scale bars, 50 μm. Genetic background, W22.

### ARF3 repression spatially patterns pectin dynamics and cell wall properties

ARF3 functions through a non-canonical auxin-response pathway to regulate gene expression^22,23^. To determine how ARF3 spatial patterning is translated into developmental patterning, we performed DNA affinity purification sequencing (DAP-seq)^24^ for three maize ARF3 paralogs (**Supplementary Fig. 2, Supplementary Dataset 1**). In total, 5,457, 8,922, and 6,001 binding regions were identified for ARF3a/ZmARF24, ARF3b/ZmARF23, and ARF3d/ZmARF12, respectively. These binding sites are significantly enriched for the auxin-responsive element TGTCGG (hypergeometric test, *P* < 1 × 10⁻¹⁰⁶⁸) and are predominantly located in gene promoters/UTRs (∼26%) and exons (∼20%), as well as within distal intergenic regions (∼30%), reflecting the large size and complex regulatory architecture of the maize genome^25^. Focusing on genes located within 10 kb of ARF3 binding sites, we identified 3,315 genes targeted by at least two ARF3 paralogs (**Supplementary Fig. 2C**).

To distinguish targeted genes regulated by ARF3 during shoot stem cell specification, we compared the transcriptomes of *lbl1-rgd1* and wild-type embryos at the early transition stage in two maize genetic backgrounds, W22 and B73 (**Supplementary Dataset 2**). This identified 106 and 1,061 differentially expressed genes (DEGs) in the W22 and B73 comparisons, respectively. Given the higher penetrance of shoot meristem-less embryos in the W22 background (**Supplementary Fig. 1**), the lower number of DEGs in W22 likely reflects the sampling of slightly earlier staged embryos, and as such may capture the earliest transcriptional changes resulting from tasiARF loss. Integration of the *lbl1-rgd1* DEGs with ARF3 DAP-seq targets revealed a total of 320 ARF3-bound genes that are differentially expressed upon loss of tasiARF-mediated ARF3 regulation, including 196 upregulated and 124 downregulated genes (**Fig. 2B**). As *Kn1* is not among these targets, we sought to investigate potential regulatory events linking ARF3 activity to stem cell specification. Gene Ontology (GO) enrichment analysis of the ARF3-targeted DEGs identified 13 significantly enriched terms, four of which are associated with shoot apical meristem function and adaxial-abaxial polarity, as expected (**Fig. 2C**). Strikingly, the remaining terms are all linked to cell wall-related pathways, pointing to a strong connection from ARF3 to cell wall remodeling during embryogenesis. Indeed, among the 14 ARF3-targeted DEGs identified at the early transition stage in W22, three genes are involved in pectin modification and degradation^26^, encoding a pectin methylesterase (*PME36*, Zm00001eb074530), a β-D-xylosidase (*BXS1*, Zm00001eb431850), and a polygalacturonase (*PGL43*, Zm00001eb359080). These genes show strong expression during the transition and coleoptile stages in wild-type embryos^27^, are downregulated in *lbl1-rgd1* embryos of B73 and W22, and are bound by one or more ARF3 paralogs at their gene bodies and proximal regulatory regions (**Fig. 2D-2E, Supplementary Fig. 3A**). Taken together, the data point to a role for the tasiARF-ARF3 regulatory module in embryonic shoot stem cell specification at least in part through regulation of genes associated with pectin dynamics and cell wall remodeling.

To verify this proposed role for ARF3, we performed whole-mount immunofluorescence using JIM7 and LM19 antibodies, which recognize highly methyl-esterified and low methyl-esterified pectins, respectively^28,29^. In wild-type embryos, JIM7 signal is broadly distributed across the adaxial surface of the proembryo, but signal intensity is consistently reduced in the presumptive shoot stem cell region of transition and coleoptile stage embryos (**Fig. 2F**). On the other hand, the LM19 signal appears enriched within the presumptive shoot stem cell domain, revealing a largely complementary pattern of pectin modification (**Supplementary Fig. 3B**). By contrast, in *lbl1-rgd1* embryos, both JIM7 and LM19 signals are perturbed and no longer display the respective spatial accumulation patterns observed in the wild-type (**Fig. 2F, Supplementary Fig. 3B**). These results indicate that ARF3 directly regulates cell wall remodeling pathways and that its local depletion in presumptive shoot stem cells permits formation of a specialized pectin modification domain required for niche specification. Ectopic ARF3 activity perturbs this spatial cell wall signature and disrupts niche formation.

### tasiARF-ARF3 generates a localized auxin minimum required for shoot stem cell fate

Pectin dynamics and cell wall remodeling are increasingly recognized as key regulators of plant morphogenesis, with mechanical properties of the cell wall known to influence the polarity, trafficking, and membrane dynamics of PIN-FORMED (PIN) auxin efflux carriers that direct auxin transport^30–35^. In addition, DAP-seq analysis showed binding of ARF3 paralogs at the *ZmPIN1* (Zm00001eb372180) and *ZmPIN2* (Zm00001eb389260) loci (**Supplementary Fig. 2D**). We therefore tested whether ARF3 spatial patterning affects PIN localization using whole-mount immunofluorescence staining with anti-ZmPIN1 and anti-ZmPIN2 antibodies (**Fig. 3A, Supplementary Fig. 4A**). In wild-type, signals for both PIN proteins are initially distributed broadly across the proembryo surface, although cellular polarity could not be resolved, likely owing to technical limitations of the immunostaining procedure. As development proceeds, PIN signals are first depleted from the presumptive shoot stem cells, generating a distinct expression minimum, and subsequently increase in the surrounding cells that give rise to the ring-shaped coleoptile primordium. In *lbl1-rgd1*, the PIN expression minimum fails to form, and no PIN-marked coleoptilar ring is established. Immunostaining with two anti-indole-3-acetic acid (IAA) antibodies recognizing distinct IAA epitopes (see Methods) revealed a comparable auxin minimum at the presumptive stem cell niche of wild-type transition-stage embryos, followed by auxin accumulation in the initiating coleoptile (**Fig. 3B, Supplementary Fig. 4B**). Importantly, no comparable localization minimum was observed in the anti-α-tubulin control (**Supplementary Fig. 4C**). In *lbl1-rgd1*, where neither *ARF3a* nor PIN forms a local expression minimum (**Fig. 2A**), IAA signals remain broadly distributed across the adaxial embryo surface (**Fig. 3B, Supplementary Fig. 4B**). tasiARF-mediated repression of ARF3 thus patterns the distribution of auxin on the adaxial face of the embryo, generating a localized auxin minimum conducive to shoot stem cell specification and a surrounding auxin maximum that drives emergence of the coleoptile.

**Figure 3.**
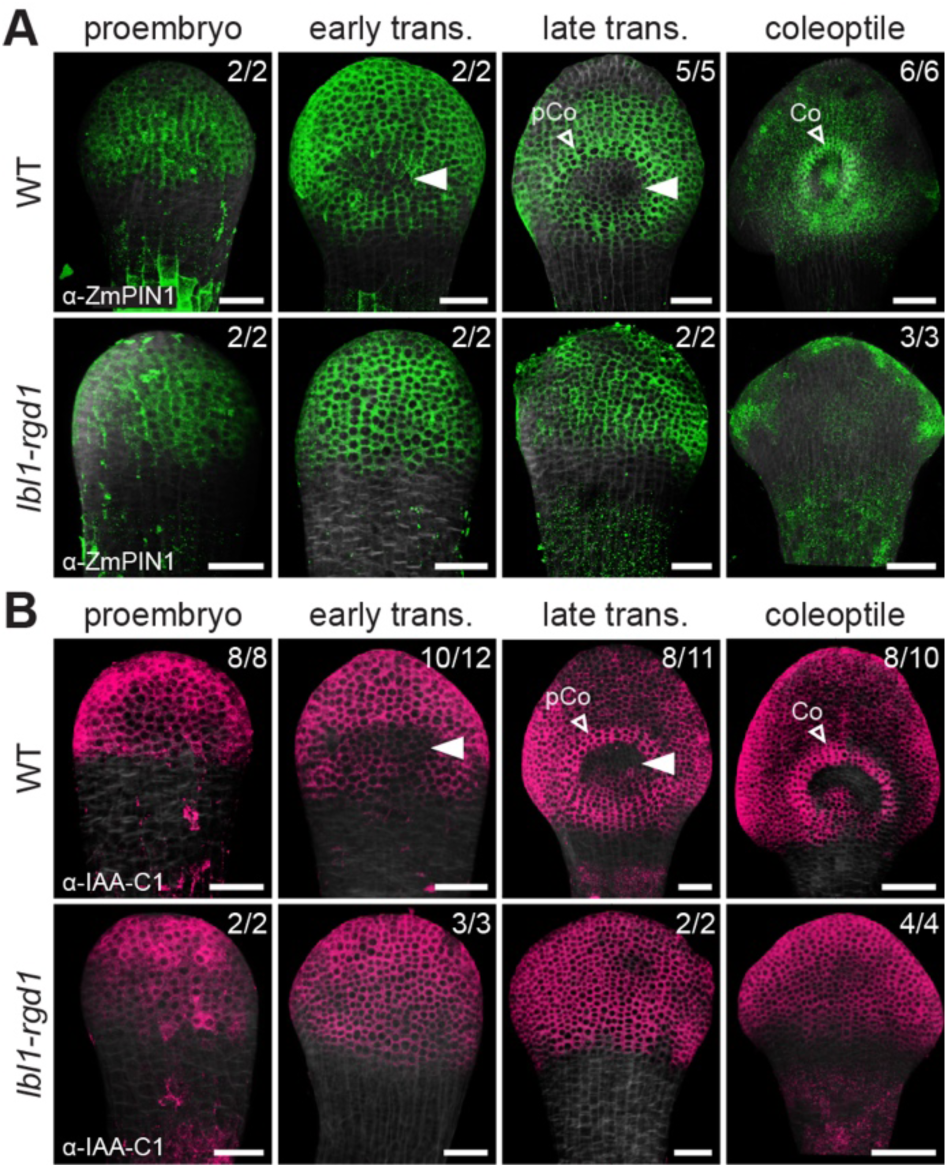
The tasiARF-ARF3 module patterns auxin transport to establish a localized auxin minimum. (**A**, **B**) Adaxial views of anti-ZmPIN1 (**A**) and anti-IAA-C1 (**B**) whole-mount immunofluorescence-stained wild-type and *lbl1-rgd1* sibling embryos show a local depletion in ZmPIN1 and auxin accumulation at the initiating shoot stem cell niche in wild-type transition stage embryos (arrowheads), not evident in *lbl1-rgd1.* Signal intensities for both antibodies become enriched in the initiating surrounding coleoptile primordium. Equivalent spatial distribution patterns are observed for anti-ZmPIN2 and anti-IAA-N1, not for anti-TUA (positive control) and Alexa Fluor 488 alone (negative control) (**Supplementary Fig. 4**). pCo, presumptive coleoptile; Co, coleoptile; trans., transition stage. Scale bars, 50 μm. Genetic background, W22.

Spatial patterns of cell wall modifications and auxin signaling can generate differential growth and mechanical conflicts between neighboring tissues that reinforce cell fate decisions^34–39^. We therefore examined patterns of cortical microtubule (CMT) alignment, which reflect the principal directions of tensile stress^37,38^. In the early transition stage, CMTs in protodermal cells of the presumptive shoot stem cell niche are arranged largely isotropically (**Supplementary Fig. 5**). As the coleoptile emerges, the arrangement of CMTs at the boundary between the stem cell domain and growing coleoptile primordium becomes increasingly anisotropic, aligning with the circumference of the developing meristem. CMTs within the shoot stem cell domain itself, however, remain largely isotropic. In *lbl1-rgd1*, CMT organization in adaxial protodermal cells remains isotropic throughout development, consistent with a failure to establish distinct stem cell and coleoptile domains (**Supplementary Fig. 5**). These findings suggest that ARF3-mediated patterning of cell wall properties and auxin signaling promotes differential growth between the embryonic shoot stem cells, the surrounding coleoptile, and intervening boundary, thereby generating inter-tissue mechanical conflicts reflected in characteristic CMT reorganization.

Collectively, these results identify the tasiARF-ARF3 module as the upstream organizer of embryonic shoot stem cell specification in maize. By spatially restricting ARF3 activity, tasiARF coordinates cell wall remodeling and PIN localization to generate a localized auxin minimum at the presumptive stem cell niche. The resulting redistribution of auxin toward surrounding cells promotes coleoptile formation, which in turn may reinforce the auxin minimum through changes in tissue mechanics associated with regional differences in growth^32–35,39^. The findings thus reveal a hierarchical morphogenic signaling cascade that couples small RNA-mediated positional information, cell wall remodeling, auxin transport, and organ growth to robust embryonic shoot stem cell specification.

### Natural variation at a QTL controlling MAP65-3 expression buffers tasiARF loss during stem cell specification

Natural genetic variation can reveal mechanisms that buffer developmental programs against perturbation. Although tasiARF is required for embryonic shoot stem cell specification in W22, where all *lbl1-rgd1* embryos fail to form a shoot apical meristem, more than half of *lbl1-rgd1* embryos in the B73 background (designated *lbl1-rgd1*^B73^) still develop a shoot system, albeit with disrupted adaxial-abaxial leaf polarity (**Fig. 4A, Supplementary Fig. 1**). Similarly, nearly half of *lbl1-rgd1*^B73^ embryos tested established an irregularly shaped auxin minimum on the adaxial embryo surface, despite persistent defects in scutellum morphology (**Fig. 4A**). These observations point to natural genetic variation that can partially compensate the loss of tasiARF-mediated patterning and buffer defects in the morphogenic cascade specifying embryonic shoot stem cell fate. To identify the underlying mechanisms, we generated F_2_ mapping populations from intercrossing between heterozygous *lbl1-rgd1*^B73^ and *lbl1-rgd1*^W22^ individuals (**Supplementary Fig. 6A**). A key advantage of using the *lbl1-rgd1* allele in this study is its tight genetic linkage to *yellow endosperm1* (*y1*), which affects carotenoid biosynthesis in the seed in a dosage-dependent manner, thus enabling the identification of homozygous and heterozygous *lbl1-rgd1* individuals based on seed color^40,41^. Using these populations, we mapped a major quantitative trait locus (QTL) to an ∼10-Mb interval on chromosome 8 that modulates the penetrance of shoot stem cell niche formation in a dosage-dependent manner (**Fig. 4B**).

**Figure 4.**
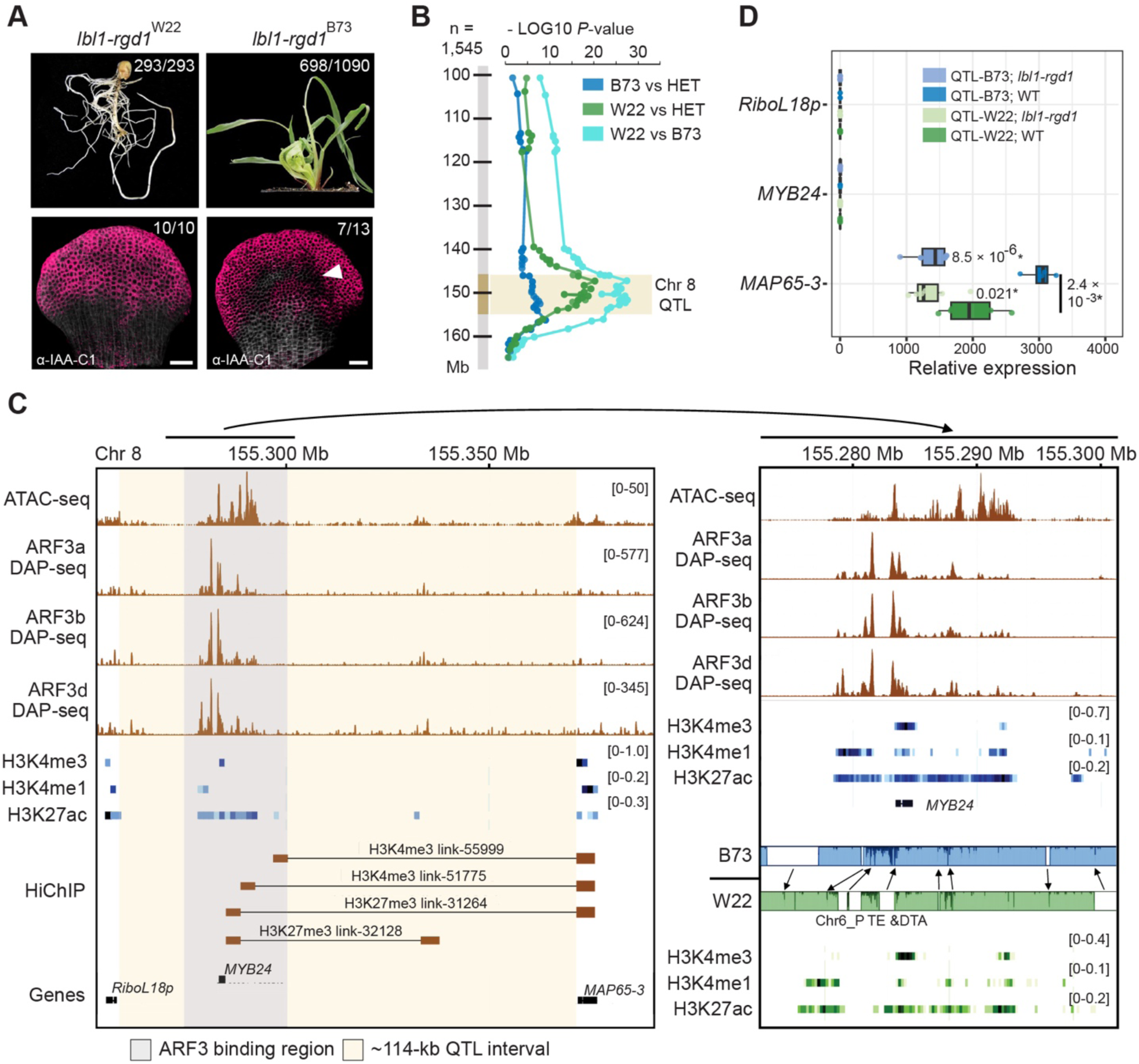
Natural variation at a QTL controlling MAP65-3 expression buffers tasiARF loss during stem cell specification. (**A**) The *lbl1-rgd1* shoot meristem-less phenotype while fully penetrant in W22, is partially suppressed in the B73 genetic background (top), coincident with formation of an irregularly shaped auxin minimum visualized by whole-mount immunofluorescence using anti-IAA-C1 antibody in *lbl1-rgd1* transition stage embryos (bottom). Scale bars, 50 μm. (**B**) High-throughput SNP genotyping of 1,545 *lbl1-rgd1* mutant progeny from (*lbl1-rgd1*^B73^/+ × *lbl1-rgd1*^W22^/+) F₂ families identifies a major QTL on chromosome 8 affecting the penetrance of the *lbl1-rgd1* shoot stem cell defects. Each dot represents an SNP marker plotted by its association with the phenotype as -log10 (*P*-value). Genomic coordinates are based on the B73 RefGen_v3 genome assembly. (**C**) Analysis of NIL-derived F2 mapping populations positions the QTL within a ∼114-kb interval encompassing *MYB24* and flanked by *MAP65-3* and *RiboL18p*. Enrichment profiles of chromatin accessibility (ATAC-seq), ARF3a/b/d DAP-seq binding, histone H3 modifications, and long-range chromatin interactions (HiChIP) across the QTL interval are shown (left). Comparison of the ∼20 kb ARF3-binding region in B73 (blue) and W22 (green) reveals genotype-specific polymorphisms and transposable element insertions associated with differences in the local distribution of active histone marks. Shaded, sequence conservation; unshaded, transposon insertions; dark shared, nucleotide sequence divergence. Genomic coordinates are based on the B73 RefGen_v5 (NAM) genome assembly. See **Supplementary Fig. 6** for more details. (**D**) Box plots showing qRT-PCR analysis of *MYB24*, *MAP65-3*, and *RiboL18p* expression in NIL^B73^-33 transition-stage embryos homozygous for either QTL-B73 or QTL-W22 (**Supplementary Fig. 6**) shows *MAP65-3* transcript levels are reduced in W22 and in *lbl1-rgd1* embryos. At least four biological replicates were analyzed. *P* values were calculated using a two-tailed Student’s *t*-test assuming unequal variance.

To fine map the QTL, we developed near isogenic lines (NILs) through marker-assisted iterative backcrossing of QTL-W22 and QTL-B73 into the B73 and W22 backgrounds, respectively. The initial 23 NILs were scored for penetrance of the shoot meristem-less phenotype in self-crossed offspring across three genotypes: homozygous QTL-B73, QTL-W22, and heterozygous QTL-B73/QTL-W22, enabling the refinement of the QTL interval. Selected NILs were further backcrossed to reduce introgression size, yielding 12 additional NILs that delimit the QTL to an ∼114-kb region containing a single annotated gene encoding a MYB family transcription factor (*MYB24*; Zm00001eb360510) (**Fig. 4C, Supplementary Fig. 6B-6C**). However, *MYB24* transcripts were scarcely detected in transition- and coleoptile-stage embryos, raising doubts about its contributions to the QTL (**Fig. 4D, Supplementary Dataset 2, Supplementary Fig. 7**). Interestingly, analysis of publicly available HiChIP datasets generated using antibodies against H3K4me3 and H3K27me3^25^ identified long-range chromatin interactions between an ∼20-kb region encompassing the transcriptionally silent *MYB24* locus and *MAP65-3* (*MICROTUBULE-ASSOCIATED PROTEIN 65-3*, also known as *INDETERMINATE GAMETOPHYTE2*^42^; Zm00001eb360520), located immediately downstream of the QTL interval (**Fig. 4C**). These interactions suggest that distal regulatory elements located within the QTL interval may control *MAP65-3* expression. Consistent with this notion, chromatin profiling reveals strong ATAC-seq signals together with enrichment of active histone marks (H3K4me1, H3K4me3, and H3K27ac)^43,44^, indicative of an open transcriptionally active chromatin configuration, specifically at both the *MAP65-3* locus and the ∼20-kb region surrounding *MYB24* (**Fig. 4C**). Notably, all three ARF3 DAP-seq datasets show prominent binding within this ∼20-kb region in B73, coinciding with chromatin accessibility and the long-range interactions to *MAP65-3* (**Fig. 4C**), suggesting ARF3-mediated long-distance regulation of *MAP65-3* expression. Comparative genomic analysis further identified multiple transposon insertion polymorphisms (including *Chr6_P*, *TE* and *DTA*) between W22 and B73 within these ARF3-binding regions, accompanied by local redistribution of active histone marks (**Fig. 4C**). In line with this, *MAP65-3* is strongly expressed in transition- and coleoptile-stage embryos, with QTL-B73 supporting higher expression than QTL-W22 in wild-type embryos in the NIL^B73^ background (**Fig. 4D, Supplementary Fig. 7**). Therefore, these observations identify *MAP65-3,* rather than *MYB24*, as the likely basis for the chromosome 8 QTL and suggest that natural variation within distal ARF3-responsive regulatory elements modulates its expression.

MAP65-3 is a microtubule-bundling protein that organizes mitotic microtubule arrays and stabilizes the phragmoplast to orient cell division and ensure proper cell plate formation during cytokinesis^45–47^. To validate its genetic contribution to the variable shoot meristem-less phenotype of *lbl1-rgd1*, we obtained three *map65-3* lesion alleles. *map65-3-ems1* and *map65-3-ems2* result from single nucleotide substitutions that affect a splice acceptor site and create a premature stop codon, respectively (**Fig. 5A**). When homozygous in the B73 genetic background, these alleles prevent both endosperm and embryo formation, generating an empty pericarp phenotype (**Supplementary Fig. 8A**), in line with the essential role for *MAP65-3* in cell division^45–47^. A third allele, *map65-3-bonnmu*, carries a *Mutator* transposon insertion in the penultimate exon encoding part of the highly conserved MAP65-ASE domain (**Fig. 5A**). Accordingly, *map65-3-bonnmu* confers a milder kernel phenotype when homozygous and in combination with *map65-3-ems1* or *map65-3-ems2* (**Supplementary Fig. 8A**). Based on the dosage-dependent effect of the chromosome 8 QTL (**Fig. 4B**), defective *map65-3* alleles paired with QTL-W22 are expected to promote penetrance of the shoot meristem-less *lbl1-rgd1* phenotype when compared to the QTL-B73/QTL-W22 combination. Indeed, pairing *map65-3-ems1* or *map65-3-ems2* with QTL-W22 in the B73 background increased the frequency of shoot meristem-less *lbl1-rgd1* embryos by approximately 17.5% (Mann-Whitney U test, *P* = 0.038), whereas the weak allele *map65-3-bonnmu* did not produce a comparable effect (**Fig. 5B**). Additionally, the weak *map65-3-bonnmu* allele in B73, which by itself does not cause a shoot meristem-less phenotype, in combination with *lbl1-rgd1*^B73^ significantly increased the frequency of shoot meristem-less *lbl1-rgd1* embryos from ∼29.3% to ∼56.4% (**Fig. 5C**; *P* = 3.9 × 10⁻⁴). Collectively, these genetic analyses demonstrate that differential regulation of *MAP65-3* expression underlies the QTL controlling the variable penetrance of the *lbl1-rgd1* shoot meristem-less phenotype in B73 versus W22.

**Figure 5.**
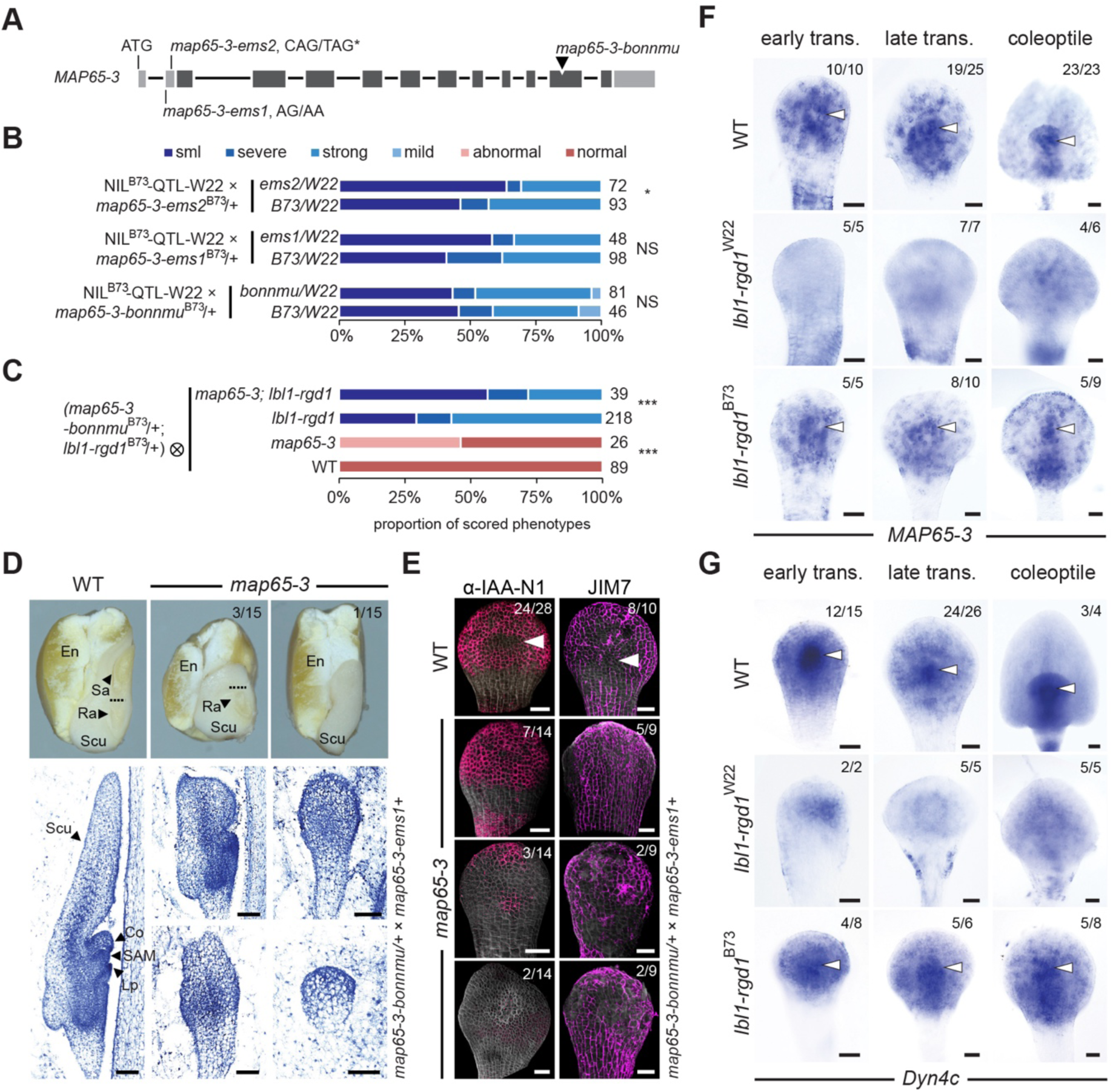
Localized *MAP65-3* expression buffers embryonic shoot stem cell specification upon tasiARF loss. (**A**) Schematic representation of *MAP65-3* with the genetic lesions of mutant alleles indicated. Boxes, coding sequences; dark shading, conserved MAP-ASE domain-encoding region. (**B**) Phenotypic severity of *lbl1-rgd1* progeny with different *MAP65-3* genotypes in the indicated crosses shows that pairing *map65-3-ems1* with QTL-W22 significantly increases the frequency of shoot meristem-less *lbl1-rgd1* embryos. Genetic background, B73. Phenotypes were classified as shoot meristem-less (sml), severe, strong, or mild, as defined in **Supplementary Fig. 1**. (**C**) Phenotypic severity of *map65-3-bonnmu*; *lbl1-rgd1* double mutants versus *lbl1-rgd1*, *map65-3-bonnmu* or wild-type segregants from the indicated F₂ population shows perturbation of *MAP65-3* enhances the penetrance of the shoot meristem-less phenotype. “abnormal” and “normal” indicate the presence or absence, respectively, of *map65-3-bonnmu*-specific seedling phenotypes. Numbers to the right of each bar indicate the total number of progeny analyzed. Statistical significance was assessed using the Mann–Whitney U test. *, *P* < 0.05; ***, P < 0.001; ns, not significant. (**D**) Longitudinal sections through mature kernels (top) or 1^st^ leaf stage embryos (paraffin sections, bottom) of wild-type (left) and *map65-3* (right) progeny from *map65-3-bonnmu*/+ × *map65-3-ems1*/+ crosses exemplify the range of abnormal *map65-3* embryo morphologies. En, endosperm; Sa, shoot apex; Ra, root apex; Scu, scutellum; Co, coleoptile; SAM, shoot apical meristem; Lp, leaf primordium. See **Supplementary Fig. 8** for more details. (**E**) Adaxial views of anti-IAA-N1 (left) and JIM7 whole-mount immunofluorescence-stained early transition stage wild-type (top) and *map65-3* (bottom) embryos shows variable and abnormal spatial distributions for auxin and highly methyl-esterified pectin in *map65-3*. Arrowhead, auxin minimum at the presumptive stem cell niche in wild-type. (**F**, **G**) Whole-mount RNA *in situ* hybridization using antisense probes against *MAP65-3* (**F**) and *Dyn4c* (***G***) shows expression associated with stem cell specification in wild-type (top), is severely reduced in *lbl1-rgd1^W22^* (middle) but variably maintained in *lbl1-rgd1^B73^*. Arrowheads, regions of enriched transcript accumulation. trans., transition stage. Scale bars, 50 μm.

### Localized MAP65-3 expression reinforces embryonic shoot stem cell specification

To investigate how MAP65-3 contributes to the morphogenetic program specifying embryonic shoot stem cell fate, we analyzed the phenotypes of partial *map65-3* loss-of -unction mutants. Combining the *map65-3-ems1* and *map65-3-bonnmu* alleles produced a spectrum of embryonic phenotypes, ranging from early embryonic arrest and loss of shoot or root formation to milder disruptions of the shoot-root axis (**Fig. 5D-5E, Supplementary Fig. 8B**). These defects are associated with perturbations in the spatial distribution of auxin, which either persists in the presumptive shoot stem cell domain or becomes dispersed and diminished across the adaxial embryo surface (**Fig. 5E, Supplementary Fig. 9A**). The pattern of methyl-esterified pectin accumulation, revealed by JIM7 immunostaining, likewise becomes variable across the embryo (**Fig. 5E**). Weaker *map65-3-bonnmu*/*map65-3-ems1* mutants, as well as plants homozygous for *map65-3-bonnmu*, further display defects in leaf patterning, affecting epidermal differentiation, sheath-blade boundary placement, mediolateral symmetry, margin formation, and leaf initiation resulting in pinhead-like shoots (**Supplementary Fig. 8B-8C**). Interestingly, these defects resemble those of auxin-transport and -signaling mutants^48–52^. As such, the findings show that MAP65-3 is required for developmental patterning throughout shoot ontogeny, presenting links between cell division orientation, cell wall placement, and auxin dynamics, including during embryonic shoot stem cell specification.

*MAP65-3* transcript levels are significantly reduced in *lbl1-rgd1* embryos irrespective of the QTL genotype (**Fig. 4D**), yet variation at this locus modulates the penetrance of the shoot meristem-less phenotype (**Fig. 5B-5C**). To resolve this seeming discrepancy, we used whole-mount RNA *in situ* hybridization to examine the pattern of *MAP65-3* expression during early embryogenesis and how this pattern changes in *lbl1-rgd1*^W22^ versus *lbl1-rgd1*^B73^ (**Fig. 5F, Supplementary Fig. 9C**). *MAP65-3* is predominantly expressed in actively dividing cells^45^. Consistent with this, the *MAP65-3* expression pattern in early developing embryos mirrors that of the cell cycle marker histone *H4* (Zm00001eb187430). Both genes show stochastic expression in transition stage embryos with enrichment in and around the stem cell precursors expressing the shoot stem cell marker *Dyn4c* (Zm00001eb011320)^53,54^. At later stages, *MAP65-3* and *H4* strongly mark both the embryonic shoot stem cell domain and the initiating coleoptile. Therefore, MAP65-3 activity is closely associated with embryonic shoot stem cell specification and subsequent coleoptile formation, in line with classical studies that defined the transition stage by the rapid proliferation and oriented divisions of cells in the stem cell precursor region^10,11^. Importantly, *Dyn4c* expression and localized expression of *MAP65-3* and *H4* at the site of the presumptive stem cell precursors are disrupted in *lbl1-rgd1*^W22^ but partially retained in *lbl1-rgd1*^B73^ embryos (**Fig. 5F-5G, Supplementary Fig. 9B**). Taken together, the observations indicate that natural variation at a distal regulatory region of *MAP65-3* underlies the QTL modulating embryonic shoot stem cell specification upon tasiARF loss. The *cis*-regulatory variation in B73 preserves a residual MAP65-3 expression domain, despite overall transcript depletion, that buffers defects in the morphogenic signaling cascade downstream of tasiARF, thereby promoting specification of the embryonic shoot stem cell niche.

## Discussion

Our study clarifies how embryonic shoot stem cells are specified during maize embryogenesis, elucidating a fundamental development process critical to the formation of virtually all aerial organs. We show that this event is driven by a hierarchical morphogenic signaling network initiated by the small RNA tasiARF in the adaxial protoderm (**Fig. 6**). By spatially restricting ARF3 expression, tasiARF remodels cell wall properties and patterns PIN-dependent auxin transport, thereby establishing a localized auxin minimum permissive for stem cell fate. This morphogenic program is interdependently coupled to cell division patterning. Auxin redistribution to cells surrounding the presumptive stem cell niche simultaneously promotes coleoptile formation, generating growth and mechanical asymmetries that may further reinforce the differential auxin distribution patterns^32–34,37,39^. In addition, natural variation at a cis-regulatory locus controlling MAP65-3 expression, which is under tasiARF-ARF3 control, buffers defects in the developmental program. The findings thus reveal an intricate morphogenic circuit that couples small RNA-mediated positional information to a self-stabilizing network interdependently linking cell wall mechanics, auxin signaling and regulated cell division, to specify the embryonic shoot stem cell niche.

**Figure 6.**
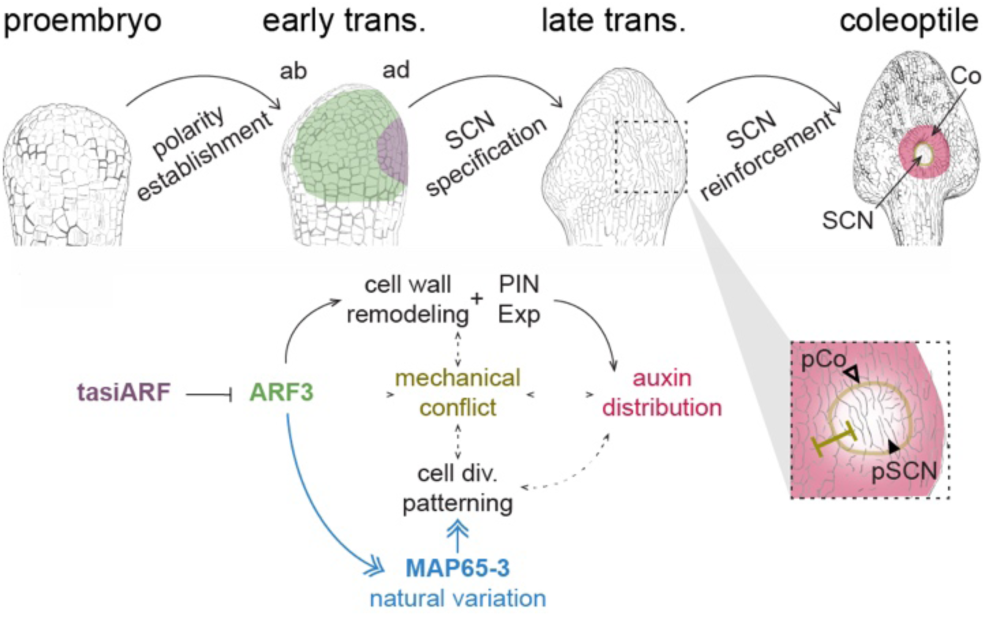
Model for the tasiARF-initiated self-stabilizing morphogenic circuit underlying embryonic shoot stem cell specification in maize. The transition from proembryo to early transition stage embryo is accompanied by acquisition of adaxial-abaxial polarity. tasiARF produced in the adaxial protoderm spatially restricts ARF3 expression, which coordinates cell wall remodeling and PIN expression, establishing a local low-auxin environment permissive for shoot stem cell specification. Auxin redistribution towards cells surrounding the presumptive shoot stem cell niche (SCN) promotes coleoptile formation, generating cell division and mechanical asymmetries between the developing coleoptile and stem cell niche that reinforce the tasiARF-initiated signaling cascade. In parallel, natural variation at an ARF3-bound regulatory region modulates MAP65-3 expression, partially restoring cell division and auxin distribution patterns upon tasiARF loss. tasiARF, ARF3, auxin, MAP65-3, and mechanical conflict are depicted in purple, green, red, blue, and gold, respectively. pCo, incipient coleoptile; pSCN, presumptive shoot stem cell niche; Co, coleoptile; SCN, shoot stem cell niche; trans., transition stage.

As in the dicot *Arabidopsis*^8,9,55,56^, embryonic shoot stem cell specification in maize requires the establishment of a localized low auxin environment permissive for stem cell fate. The conservation of this developmental output is notable given the profound differences in embryonic architecture between monocots and dicots, and suggests that the requirement for a low-auxin niche is a conserved principle of shoot stem cell specification across flowering plants. The upstream mechanisms generating this minimum, however, appear to have diverged in accordance with embryonic architecture. In Arabidopsis, the auxin minimum forms between two cotyledon-associated auxin maxima during the establishment of bilateral symmetry^8,9,55^. The underlying mechanism remains unclear but conceivably proceeds directly as a consequence of auxin drainage toward the developing cotyledon primordia. In monocots, the single cotyledon or scutellum occupies the side of the embryo opposite the future stem cell niche and therefore cannot establish an equivalent drainage-based auxin minimum^9^. Instead, tasiARF has been co-opted as an upstream positional signal that initiates the morphogenic circuit culminating in the formation of a localized auxin minimum, a role that is not evident in dicots. Our findings thus reveal lineage-specific innovation to establish a conserved stem cell fate-permissive low auxin environment within the divergent embryonic architecture of monocotyledonous cereals.

Our study also sheds light on the developmental origin and function of the coleoptile during monocot embryogenesis, supporting the view that it arises directly from proembryonic cells^57–59^, in conjunction with shoot stem cell niche specification. As the auxin minimum is established within the presumptive stem cell domain, auxin accumulates in surrounding cells giving rise to the coleoptile primordium. Continued auxin transport toward the developing coleoptile primordium raises the intriguing possibility that this assumes a function analogous to the Arabidopsis cotyledons, serving as a local auxin sink, but one that reinforces auxin depletion from the establishing stem cell niche. Moreover, coleoptile emergence is accompanied by differential growth relative to the niche, generating inter-tissue mechanical conflicts that may further stabilize local auxin distribution patterns^32–34,37,39^. The coleoptile therefore appears to arise as a consequence of the tasiARF-initiated morphogenic program while subsequently serving as a reinforcing element that amplifies this program’s output.

Because embryonic shoot stem cell specification is indispensable for postembryonic development, the underlying morphogenetic program would be expected to be highly robust. Surprisingly, the core components of tasiARF biogenesis - LBL1, RDR6, RGD2/AGO7, and DCL4 - are each encoded by a single gene in the maize genome^19,20^, indicating that developmental robustness cannot be achieved through genetic redundancy within the pathway itself. Our characterization of a QTL modulating the penetrance of the shoot meristem-less phenotype in *lbl1-rgd1* reveals that the tasiARF-initiated morphogenic program in certain genetic backgrounds is buffered by differential expression of MAP65-3. Natural variation at an ARF3-bound distal regulatory region variably preserves MAP65-3 expression in the absence of tasiARF, partially restoring the auxin distribution patterns required for stem cell specification, while genetic perturbation of *MAP65-3* enhances the penetrance of the shoot meristem-less phenotype. This links enriched expression of MAP65-3 in and around the stem cell precursors to their specification. As a key determinant of cell division orientation and cell plate formation^45–47^, MAP65-3 may contribute to a mechanical environment permissive for shoot stem cell specification^30,32–34,37^. Robustness in embryonic shoot stem cell specification is thus attained not through redundancy within the tasiARF pathway, but likely through reciprocal interactions between regulated cell division, mechanical patterning, and auxin transport that interdependently reinforce the tasiARF-initiated morphogenic cascade.

Overall, our study establishes a mechanistic framework for shoot stem cell specification during monocot embryogenesis. This reveals a conserved developmental principle, the requirement for an auxin-depleted environment, that is shared between dicots and monocots. However, the upstream mechanisms generating this environment have diverged to accommodate fundamentally different embryonic architectures. In monocots, this conserved developmental outcome is achieved through lineage-specific redeployment of the deeply conserved tasiARF-ARF3 module^60,61^, which functions as a transcriptional regulator of the mechanical and auxin signaling environment, into a new developmental context. Our findings thus illustrate how evolution can preserve a core developmental principle while rewiring the regulatory network that implements it. An intriguing question for future study is how the proembryo first acquires adaxial-abaxial polarity, and whether the tasiARF pathway also plays a role therein. More broadly, given the striking parallels between embryonic stem cell specification and *de novo* shoot regeneration^6,7^, this work identifies molecular entry points for engineering embryogenic competence and regeneration capacity, providing new avenues for the improvement of cereal crops.

## Methods

### Plant materials and cultivation

The maize tasiRNA biogenesis mutant alleles *lbl1-rgd1* and *dcl4-2* have been described previously^19,20^. These alleles were introgressed into W22 and B73 inbred backgrounds for at least four generations prior to phenotypic analysis. Mutants were identified based on phenotypic segregation in selfed heterozygous progeny, with *lbl1-rgd1* additionally inferred based on its linkage to the *y1* mutant allele^41^. The *map65-3-ems1* and *map65-3-ems2* alleles were generated by EMS mutagenesis (maizeems.qlnu.edu.cn) in a B73 background and gifted by Yongrui Wu (CEMPS, CAS). The *map65-3-bonnmu* allele was identified from the BonnMu project (www.inres.uni-bonn.de/cfg/en/c-f-g/research/bonnmu)^62,63^ and was introgressed into B73 for at least three generations. Genotyping primers for the *map65-3-ems1/-ems2/-bonnmu* and *lbl1-rgd1* alleles are listed (**Supplementary Dataset 3**). Plant cultivation and genetic crosses were performed in the field or under controlled conditions in growth chambers. Here, plants were grown under long-day conditions (16 h light/8 h dark) at 25–28 °C during the day and 20–22 °C at night, with relative humidity maintained at ∼50–60%.

### Morphological analysis

Multiple stages of early maize embryos were dissected from pollinated ears at 6-8 days after pollination (DAP), depending on growth conditions. For histological analysis, embryos were fixed in 4% paraformaldehyde (PFA) in 1× PBS-T and processed for paraffin sectioning or whole-mount staining^64,65^. For paraffin embedding, samples were dehydrated through an ethanol series, cleared in xylene, and infiltrated with Paraplast Plus (Sigma-Aldrich, Cat# P3683) using a Leica EG1140H embedding station. Sections (10 μm) were prepared using a rotary microtome (Leica RM2255), mounted on slides, deparaffinized in xylene, and rehydrated through a descending ethanol series. Sections were stained with 0.05% (w/v) toluidine blue O, rinsed in water, dehydrated, mounted, and imaged using a bright-field microscope (Zeiss Axio Imager M2). For whole-mount observation, the PFA-fixed samples were cleared in ClearSee solution (10% [w/v] xylitol, 15% [w/v] sodium deoxycholate, and 25% [w/v] urea) at 4 °C prior to staining. Samples were then stained with Fluorescent Brightener 28 (FB28; Sigma-Aldrich, Cat# F3543; 0.1% w/v), mounted in water on concave slides, and imaged using a Leica TCS SP8 confocal microscope. FB28 fluorescence was excited at 405 nm and detected at 420–480 nm. Z-stack images were processed in Fiji (v2.16.0/1.54p) and visualized using MorphoGraphX (v2.0.3).

### Paraffin-based and whole-mount RNA *in situ* hybridization

DIG-labeled RNA probes for *in situ* hybridization we generated as previously described^64^. Briefly, gene-specific coding region fragments were PCR amplified using DreamTaq DNA Polymerase with 10× DreamTaq Green Buffer (Thermo Fisher Scientific, Cat# EP0701 and B71) and cloned into pSPT18 (DIG RNA Labeling Kit, Roche, Cat# 11175025910) or pCRII-TOPO TA (Thermo Fisher Scientific, Cat# K460001). Gene inserts were amplified using pBRrevBam/pGEX_3 or M13F/M13R primers, purified by ethanol precipitation, and probes synthesized by in vitro transcription using the DIG RNA Labeling Kit (Roche, Cat# 11175025910). Probe quality was assessed by agarose gel electrophoresis and western blot analysis, and probes were partially hydrolyzed when necessary to facilitate tissue penetration, as described previously^64^. For detection of tasiARF^16,18^, a DIG-labeled locked nucleic acid (LNA) probe complementary to the mature tasiARF sequence was used (Qiagen, formerly Exiqon), with hybridization performed at 37 °C. Primer sequences used for probe synthesis are listed in **Supplementary Dataset 3**.

Paraffin-based RNA *in situ* hybridization was performed largely as previously described^64^. Briefly, maize kernels were fixed in 4% paraformaldehyde (PFA), dehydrated through a graded ethanol series, cleared in xylene, and embedded in Paraplast Plus (Sigma-Aldrich, Cat# P3683). Sections (10 μm thick) were prepared using a rotary microtome (Leica RM2255), mounted on positively charged Superfrost Plus slides (Thermo Fisher Scientific), and dried overnight at 37 °C on a histology slide heater (VWR, W10). Sections were deparaffinized, rehydrated, treated with protease, refixed, acetylated, and dehydrated prior to hybridization. Hybridizations were carried out overnight at 55 °C using DIG-labeled probes at empirically optimized concentrations (typically ∼100 ng/mL) in hybridization buffer containing 50% deionized formamide, *in situ* hybridization salts (0.375 M NaCl, 12.5 mM Tris-HCl pH 8.0, 12.5 mM sodium phosphate pH 6.8, and 6.25 mM EDTA), 12.5% dextran sulfate, 1.25× Denhardt’s solution (Sigma-Aldrich, Cat# D2532), and 1.25 mg/mL tRNA (Sigma-Aldrich, Cat# R5636). Following hybridization, samples were washed under high-stringency conditions using a graded SSC series (2×, 1×, and 0.5× SSC; Sigma-Aldrich, Cat# S6639), incubated with an alkaline phosphatase-conjugated anti-DIG antibody (Roche, Cat# 11093274910), and signals were developed using NBT/BCIP (Roche, Cat# 11681451001). The colorimetric reaction was stopped by rinsing the samples in ethanol, followed by dehydration. Slides were subsequently dehydrated, mounted, and imaged using a bright-field microscope (Zeiss Axio Imager M2).

For whole-mount RNA *in situ* hybridizations, dissected tissues were fixed in FAA (50% [v/v] ethanol, 3.7% [v/v] formaldehyde, and 5% [v/v] acetic acid), dehydrated through an ethanol series, and incubated in methanol at −20 °C. Samples were permeabilized by xylene/ethanol treatment, rehydrated, and subjected to partial cell wall digestion for 10 min at room temperature in 1× PBS-T containing 0.1% (w/v) Macerozyme R-10, 0.1% (w/v) Cellulase Onozuka RS, 0.05% (w/v) Pectolyase Y-23, 0.15% (w/v) Pectinase, and 0.1% (w/v) Hemicellulase. Samples were then treated with Proteinase K (80 μg/mL; Roche, Cat# 03115879001) for 15 min at 37 °C. After each digestion step, samples were refixed in 4% formaldehyde. Hybridization was performed overnight at 50 °C. Whole-mount samples were subsequently processed as described above and cleared in ClearSee solution prior to imaging with a bright-field microscope (Zeiss Axio Imager M2).

### Whole-mount immunofluorescence analysis

Whole-mount immunolocalization was performed as described previously^65^, with modifications depending on antibody and signal characteristics. The following primary antibodies were used: anti-ZmPIN1 (AS22 4801, Agrisera), anti-ZmPIN2 (AS21 4697, Agrisera), anti-IAA-C1 (AS09 445, Agrisera; raised against IAA conjugated via the C1 carboxyl group), anti-IAA-N1 (AS09 421, Agrisera; raised against IAA conjugated via the N1 position of indole), and anti-TUA (Sigma-Aldrich, Cat# T6074). Anti-TUA served as a positive control, whereas samples processed without primary antibody served as negative controls.

Dissected immature kernels were fixed in FAA, dehydrated through a graded ethanol series, and incubated in methanol at −20 °C. Samples were permeabilized using xylene/ethanol treatment and rehydrated. Partial cell wall digestion was performed using the enzyme mixture described for whole-mount RNA *in situ* hybridization, followed by refixation in 4% paraformaldehyde (PFA). Samples were then subjected to a mild Proteinase K treatment (20 μg/mL for 10 min at room temperature), followed by an additional refixation step in 4% PFA to enhance antigen accessibility and antibody penetration while preserving tissue structure. Samples were blocked in 1× PBS containing 3% BSA, incubated with primary antibodies (1:50-1:200 or according to the manufacturer’s recommendations) in 1× PBS containing 1% BSA overnight at 4 °C with gentle agitation, and washed extensively in 1× PBS-T. Fluorophore-conjugated secondary antibodies, including Alexa Fluor 488 anti-rabbit IgG (Invitrogen, Cat# A-11008) and anti-mouse IgG (Invitrogen, Cat# A-10680), were applied (1:100-1:400) overnight at 4 °C, followed by washing in 1× PBS-T. Samples were counterstained with FB28, mounted in water on concave slides, and imaged using a Leica TCS SP8 confocal microscope. FB28 fluorescence was excited at 405 nm and detected between 420–480 nm. Alexa Fluor 488 signals were excited at 488 nm and detected between 500–550 nm. Z-stack images were processed in Fiji (v2.16.0/1.54p) and visualized using MorphoGraphX (v2.0.3).

For JIM7 (Agrisera, AS07 023) and LM19 (Agrisera, AS11 174), the procedure was identical except that the cell wall digestion step was omitted to preserve cell wall structures. Primary antibodies were used at a 1:10 dilution, and detection was performed using Alexa Fluor 488-conjugated anti-rat IgG (Invitrogen, Cat# A-11006).

For cortical microtubule visualization, tissues were fixed in microtubule-stabilizing buffer (MTSB; 10 mM PIPES, 5 mM EGTA, and 5 mM MgSO₄, pH 6.9) containing 4% paraformaldehyde, 0.5% glutaraldehyde, 0.3% Tween-20, and 0.3% Triton X-100, followed by vacuum infiltration and fixation at room temperature. After rinsing in MTSB, samples were treated with the same enzyme mixture as above for 20-40 min depending on tissue stage, incubated with anti-TUA antibody (1:50) overnight at 4 °C, washed in 1× PBS-T, and detected with Alexa Fluor 488-conjugated anti-mouse IgG overnight at 4 °C. Confocal Z-stacks were processed in Fiji (v2.16.0/1.54p), and cortical microtubule organization at the cell surface was visualized using surface rendering. Quantification of microtubule orientation and anisotropy was performed using the FibrilTool plugin, based on manually defined cell regions^66^.

### DAP-seq profiling and analysis

Libraries were constructed from purified genomic DNA extracted from aerial tissues of 14-day-old B73 seedlings as previously described^24,67^. Briefly, 5 μg of genomic DNA was sheared to ∼200 bp using a Covaris S2 focused ultrasonicator, followed by end repair, A-tailing, and ligation to truncated Illumina adapters using the NEBNext Ultra II DNA Library Prep Kit (New England Biolabs, Cat# E7645). Adapter-ligated libraries were purified using AMPure XP beads (Beckman Coulter, Cat# A63881) and quantified using the Qubit dsDNA HS Assay Kit (Thermo Fisher Scientific, Cat# Q32854). ARF3a/ZmARF24/Zm00001eb295830, ARF3b/ZmARF23/Zm00001eb292830, and ARF3d/ZmARF12/Zm00001eb157270 transcription factor clones were obtained from the maize TF collection (grassius.org/tfomecollection). pENTR clones were recombined into the pDEST15 (ZmARF23 and ZmARF24) or pIX-HALO (ZmARF12) expression vector as described previously^24,44^. ARF23 and ARF24 were expressed in *E. coli* and purified as before^44^. To prepare the protein-bound beads, ∼20 μl or ∼5 μg purified GST-tagged ARF protein was bound to 25 μl of MagneGST beads (Promega) and incubated for 1 hour at RT. ARF12 protein was expressed in vitro using the rabbit reticulocyte TNT SP6 Coupled Transcription/Translation System (Promega, Cat# L4600) complemented with 1μl Met and 1μl Leu, according to the manufacturer’s instructions. HALO-tagged ARF12 protein was immobilized on MagneHALO magnetic beads (Promega, Cat# G7281) by incubating for 1 hour at RT. Protein-bound beads were washed and incubated with 1 μg of the adapter-ligated genomic DNA library in 1x PBS for 1 hour at RT. After six washes with 1x PBS, bound DNA was eluted in 20 μl 10 mM Tris-HCl pH 8.0, PCR-amplified with unique barcodes, cleaned on AmpureX beads, and sequenced as paired-end 150 bp reads on an Illumina NovaSeq 6000 platform. Empty pDEST15 and pIX-HALO vector samples were processed in parallel as a background control.

Sequencing reads were trimmed using Trimmomatic (v0.39) and aligned to the maize B73 RefGen_v3 (AGPv3) genome using Bowtie2 (v2.5.1). Those reads with mapping quality (MAPQ) > 30 were retained for downstream analyses. Previously published input DNA controls were processed in parallel and used for background normalization^44^. Peaks were identified using GEM (v3.4) with a stringent q-value cutoff of 1 × 10⁻⁵. Genome-wide signal tracks (bigwig files) were generated using the bamCoverage utility from deepTools2 and visualized using JBrowse 2 (v2.6.0) and the JBrowse instance embedded in MaizeGDB (jbrowse.maizegdb.org). Target genes were assigned using ChIPseeker (v1.34.1) based on proximity to annotated gene features, with promoter regions defined as 10 kb upstream of transcription start sites. For motif analysis, sequences ±100 bp around peak summits were extracted from the top 1000 peaks and subjected to *de novo* motif discovery using MEME-ChIP (v5.5.0). Identified motifs were compared with known ARF binding motifs (AuxRE), and motif enrichment and positional distribution were analyzed based on high-confidence binding sites.

### RNA-seq profiling and analysis

*lbl1-rgd1*/+ plants, introgressed into the B73 and W22 backgrounds for at least four generations, were selfed and transition-stage embryos collected at 7 days after pollination. Genotypes of the collected embryos was determined by PCR analysis of the corresponding endosperm tissue (**Supplementary Dataset 3**). Wild-type and homozygous *lbl1-rgd1* embryos were grouped into biological duplicates. Total RNA was extracted using the PicoPure RNA Isolation Kit (Arcturus, Cat# KIT0204) according to the manufacturer’s instructions, followed by DNase I treatment (Qiagen, Cat# 79254) and amplification with the TargetAmp 2-Round aRNA Amplification Kit 2.0 (Epicentre Biotechnologies, Cat# TA117). RNA quality and yield were assessed using a NanoDrop 2000 spectrophotometer and an RNA Bioanalyzer chip (Agilent RNA 6000 Nano Kit, Cat# 5067-1511). Single-end RNA-seq libraries were then constructed using standard Illumina protocols. Library quality and size distribution were assessed using a High Sensitivity DNA Bioanalyzer chip (Agilent, Cat# 5067-4626), and libraries were quantified using the KAPA Library Quantification Kit for Illumina (Roche, Cat# KK4824). Sequencing was performed on an Illumina HiSeq 2000 platform to generate 100-bp single-end reads.

Sequencing reads were aligned to the B73 RefGen_v3 (AGPv3) reference genome using STAR (v2.7.10a), with gene models defined based on the corresponding filtered gene set (FGS) annotation. Gene-level read counts were obtained from uniquely mapped reads using featureCounts (v2.0.1). Differential expression analysis was performed using DESeq2 (v1.34.0), with differentially expressed genes defined as |log2 fold change| > 1 and adjusted *P* < 0.05. Gene model conversion between the B73 RefGen_v3 (AGPv3), RefGen_v4 (AGPv4), and RefGen_v5 annotations was performed using tools available at MaizeGDB (www.maizegdb.org/gene_center/gene). Gene annotation information was obtained from NCBI Gene (www.ncbi.nlm.nih.gov/gene) and UniProt (www.uniprot.org). Gene Ontology (GO) enrichment analysis was performed using g:Profiler (biit.cs.ut.ee/gprofiler) with default settings.

### Quantitative genetic analysis and QTL mapping

For primary QTL mapping, approximately 1,600 *lbl1-rgd1* mutant F₂ progeny from the *lbl1-rgd1*^B73^/+ × *lbl1-rgd1*^W22^/+ intercross were identified by their white endosperm color, reflecting the close genetic linkage of *lbl1-rgd1* to *y1*. A portion of the endosperm from each kernel was used for DNA extraction and genotyping, while the remaining embryo-containing seed was germinated for phenotypic evaluation. Phenotypes were classified as ragged seedling or shoot meristem-less. High-throughput SNP genotyping was performed using a maize SNP array (Pioneer Hi-Bred, DuPont). Marker-trait associations were assessed using a general linear model (GLM) implemented in R, and *P*-values were calculated for each SNP based on the B73 RefGen_v3 reference genome.

For QTL fine-mapping, reciprocal near-isogenic lines (NILs) were generated by repeated (>4x) backcrossing of the chromosome 8 B73 and W22 QTL into W22 and B73, respectively. NILs with B73-W22 recombination breakpoints within the QTL interval were selfed to generate families homozygous B73, homozygous W22, or heterozygous B73-W22 across subregions of the QTL interval that segregate *lbl1-rgd1*. White *lbl1-rgd1* seed were scored and penetrance of the shoot meristem-less phenotype correlated with the chromosome 8 genotype to refine the QTL position. Select NILs were further backcrossed to identify additional informative recombination events to delimit the final 114-kb QTL interval. Molecular markers with corresponding primer sequences are provided in **Supplementary Dataset 3**.

### RT-qPCR expression analysis

*lbl1-rgd1* mutant embryos, 7-8 days after pollination, were collected from self-crosses of NIL^B73^-33-derived families homozygous for either QTL-B73 or QTL-W22. Total RNA was extracted using the Spectrum Plant Total RNA Kit (Sigma-Aldrich, Cat# STRN50), with at least four biological replicates per genotype. First-strand cDNA synthesis was performed using the iScript cDNA Synthesis Kit (Bio-Rad, Cat# 1708891). Quantitative real-time PCR (RT-qPCR) was conducted using the Luna Universal qPCR Master Mix (New England Biolabs, Cat# M3003) on a CFX96 Real-Time PCR Detection System (Bio-Rad), and data were analyzed using the associated Bio-Rad CFX Manager software. Statistical significance was assessed using a two-tailed Student’s *t*-test assuming unequal variance. Primers used for Real-Time PCR are listed in **Supplementary Dataset 3**.

### Chromatin accessibility analysis

ATAC-seq data for maize seedling leaves was retrieved from the JBrowse instance embedded in MaizeGDB (jbrowse.maizegdb.org). HiChIP data for maize seedling leaves was obtained from the Plant Epigenome Browser (epigenome.genetics.uga.edu/PlantEpigenome) and originally map positions from B73 RefGen_v4 converted to corresponding B73 RefGen_v5 positions using MaizeGDB. Datasets profiling the distribution of H3K4me3, H3K4me1, and H3K27ac in the coleoptile node in both B73 and W22 were obtained from the Gramene Maize PanGenome project (maize-pangenome.gramene.org), with correction of the control-normalized mapping data kindly provided by the authors at Cold Spring Harbor Laboratory. Data were visualized using the genome browser instance embedded within Gramene. Sequences comparisons of interval sequences between B73 and W22 were performed in Geneious Prime (v2025.0.03) using the embedded whole-genome alignment plugin based on Mauve (www.geneious.com/plugins/mauve). Sequence and annotation information were obtained from the MaizeGDB JBrowse instance.

### Genetic validation of QTL cloning

The following approaches were used to validate MAP65-3 as basis for the chromosome 8 QTL. (1) The *map65-3-ems1*, *map65-3-ems2* and *map65-3-bonnmu* alleles backcrossed for at least three generations into B73 were crossed to *lbl1-rgd1*^B73^. Resulting *map65-3/+; lbl1-rgd1/+* progeny were subsequently crossed to NIL^B73^-33, homozygous for the QTL-W22 interval and segregating for *lbl1-rgd1*. White kernels, corresponding to homozygous *lbl1-rgd1* individuals, were selected for phenotypic and genotypic analysis. Phenotypes were classified as shoot meristem-less (sml), severe, strong, or mild, as in **Supplementary Fig. 1**. Genotyping of the respective *map65-3-ems1/ems2/bonnmu* mutant allele was performed using oligonucleotide primers listed in **Supplementary Dataset 3**. (2) F₂ populations segregating for *map65-3-bonnmu* and *lbl1-rgd1* in B73 were scored for the above phenotypic classes.

## Acknowledgements

We thank members of the Timmermans lab and colleagues at the Center for Plant Molecular Biology, University of Tübingen, for helpful suggestions and comments throughout this study; Gert Huber, Philipp Becker, and Manuel Hipp for plant care; Katherine Petsch and Kevin Simcox (DuPont Pioneer Hi-Bred) for help with generating NILs and early QTL mapping; Jianming Yu and Xianran Li (Iowa State University) for QTL analysis; Martin Bayer, Sandra Richter (ZMBP - Microscopy) Agata Burian (University of Silesia in Katowice), and Xixi Zheng (University of Regensburg) for advice on confocal and bright-field microscopy; Pin-Jou Wu and Pavel Solansky for help with bioinformatic analyses; Joanna Porankiewicz-Asplund (Agrisera) for providing antibodies; Sharon Wei and Jonathan Cahn (Cold Spring Harbor Laboratory) for sharing histone modification datasets prior to publication; and Yongrui Wu (Center for Excellence in Molecular Plant Sciences, CAS) for providing the *map65-3-ems* alleles. This work was supported by the Alexander von Humboldt Professorship to M.C.P.T., by the Deutsche Forschungsgemeinschaft (DFG TI 864/5-1) through Research Unit 5235 (CSCS: Cereal Stem Cell Systems) to M.C.P.T., and by Alexander von Humboldt Research Fellowships to Q.L. and C.L.M.. A.G. and M.G. acknowledge funding from the National Science Foundation (IOS#1916804).

## Author contributions

M.C.P.T. conceived and supervised the research. Q.L. performed most experiments. C.L.M. assisted with immunofluorescence and MorphoGraphX visualization. M.C.P.T., A.F., S.K. cloned the QTL and Q.L. genetically validated its cloning. M.J. performed paraffin-based RNA *in situ* hybridizations and RNA-seq analysis. A.H. assisted with microtubule and MAP65-3 analyses. M.G. and A.G. performed DAP-seq profiling and analysis. C.M. and F.H. generated and identified the *map65-3-bonnmu* allele. Q.L., C.L.M., A.H., and M.C.P.T. interpreted the data, prepared the figures, and wrote the manuscript with input from the other authors.

## Data availability

RNA-seq data generated in this study are available from the NCBI BioProject database (www.ncbi.nlm.nih.gov/bioproject) under accession number PRJNA1465330, and ARF3a/b/d DAP-seq data are available under accession number GSE343119.

